# Analysis of Urinary Proteome Modifications in Patients with Different Glycated Hemoglobin A1c Levels

**DOI:** 10.64898/2025.12.30.697124

**Authors:** Yuzhen Chen, Youhe Gao

**Affiliations:** Gene Engineering Drug and Biotechnology Beijing Key Laboratory, College of Life Sciences, Beijing Normal University, Beijing 100875, China

**Keywords:** urine, proteomics, modifications, glycated hemoglobin A1c, diabetes mellitus, biomarker

## Abstract

Diabetes, a major global public health concern, requires early diagnosis and timely intervention. Glycated hemoglobin A1c (HbA1c) serves as a biomarker of glycemic management, with its levels showing a continuous relationship with the risk of developing diabetes. In this study, urinary proteome modifications were compared between each of the two patient groups with different HbA1c levels ([6.4±0.7]% and [8.6±1.6]%) and healthy controls. A total of 1 954 and 5 545 differentially modified peptides were identified in the two groups, respectively. Within each group, differentially modified peptides exhibiting changes from presence to absence or vice versa accounted for 48.8% and 86.5%, respectively. Additionally, results from the randomized grouping test indicated that at least 90.6% and 94.1% of these differentially modified peptides in each group were not randomly generated. In conclusion, urinary proteome modifications comprehensively and systematically reflect changes associated with elevated HbA1c levels, with distinct modification profiles corresponding to different HbA1c levels. These findings suggest that urinary proteome modifications have the potential to reflect HbA1c levels and offer a new perspective for research on the early diagnosis of diabetes.

## 1 Introduction

Diabetes is a major global public health issue. It is estimated that in 2021, the prevalence of diabetes among individuals aged 20-79 worldwide reached 10.5% (536.6 million people) and is projected to rise to 12.2% (783.2 million people) by 2045 [1]. Studies have shown that, compared with non-diabetic populations, people with diabetes face significantly increased risks of mortality from conditions such as cardiovascular disease, cancer, chronic obstructive pulmonary disease, and other diseases, which severely affect their quality of life [2]. Therefore, early diagnosis and timely intervention for diabetes are crucial.

Biomarkers are measurable changes associated with physiological or pathophysiological processes in the body, playing an important role in clinical diagnosis, treatment and prognosis. Glycated hemoglobin A1c (HbA1c) reflects the average blood glucose level over the previous 2-3 months and is used as a biomarker for diabetes [3]. The International Expert Committee recommends using HbA1c ≥ 6.5% as the diagnostic criterion for diabetes [4]. In addition, the risk of developing diabetes shows a continuous distribution based on HbA1c levels. As HbA1c approaches the diagnostic threshold for diabetes, the risk of progression to diabetes gradually increases. Among these, individuals with HbA1c ≥ 6% but < 6.5% may represent the subgroup at the highest risk of diabetes progression [4].

Urine, as a filtrate of the blood, does not need or possess homeostatic mechanisms to be stable. It accommodates and accumulates more changes without harming the body, reflecting changes in all organs and systems of the body earlier and more sensitively. Therefore, urine represents an excellent source of biomarkers [5]. Given the advantage of urine in comprehensively, systematically, and sensitively reflecting the body’s state, can the differences among patients with different HbA1c levels be characterized through urinary proteome modifications? In this study, the differences in urinary proteome modifications among patients with different HbA1c levels were explored, aiming to provide a new perspective for research on the early diagnosis of diabetes.

## 2 Materials and Methods

### 2.1 Urine Sample Information and Mass Spectrometry Detection Parameters

The mass spectrometry data used in this study were obtained from three previously published studies [6-8]. In two of these studies, patient groups exhibited different HbA1c levels: one group had an HbA1c level of (6.4±0.7)%, and the other had an HbA1c level of (8.6±1.6)%. The sample data from each patient group were separately compared with the healthy control group, respectively, and were designated as the Group A comparison and the Group B comparison. Mass spectrometry data for all samples were acquired using data-dependent acquisition (DDA) mode. Detailed sample information and mass spectrometry detection parameters are shown in Table 1.

**Table 1.**
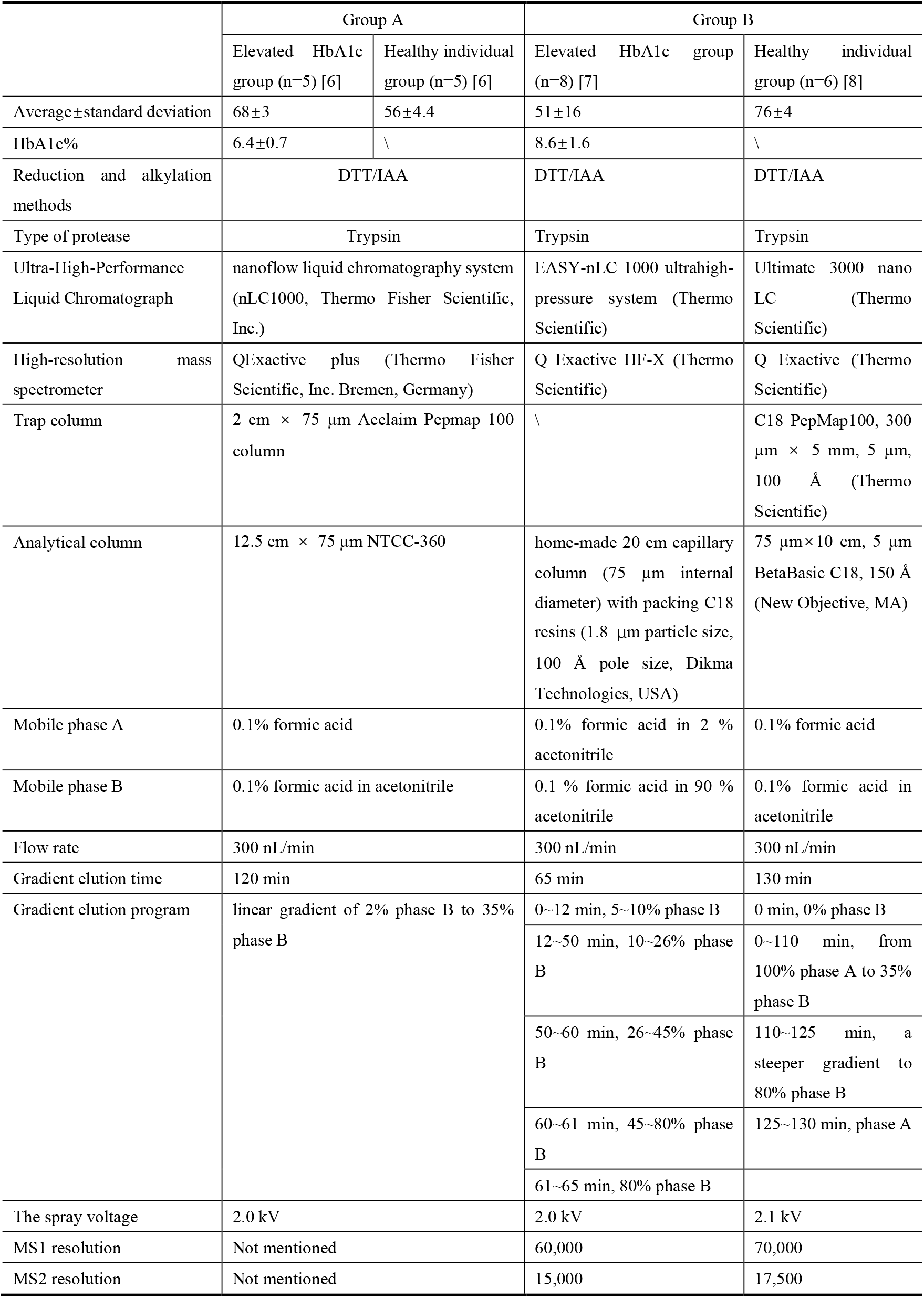
Sample information, processing methods, and mass spectrometry detection parameters.

### 2.2 Database Searching and Data Processing

Proteome modification information was obtained using pFind Studio software (version 3.2.2, Institute of Computing Technology, Chinese Academy of Sciences, Beijing, China). Label-free quantitative analysis was performed on the mass spectrometry data, with the original data files searched against the *Homo sapiens* UniProt canonical database (updated in July 2025). For the search, “HCD-FTMS” was selected for “MS Instrument”, “Trypsin_P KR P C” was chosen for trypsin digestion, and the maximum number of missed cleavages allowed per peptide was two. Both precursor and fragment tolerances were set to ±20 ppm. To identify global modifications, the “Open Search” option was selected. False discovery rates (FDRs) at the spectra, peptide, and protein levels were all kept below 1%. The number of peptide mass spectra (Total_spec_num@pep) in each sample was extracted from the analysis results in pFind Studio using the script “pFind_protein_contrast_script.py” [9, 10].

### 2.3 Data Analysis

The numbers of modified peptide mass spectra identified in the two patient groups with different HbA1c levels and in the healthy controls were compared. Differentially modified peptides were screened according to the following criteria: fold change (FC) ≥1.5 or ≤ 0.67, and *p* < 0.05 by two-tailed unpaired *t*-test analysis. Hierarchical cluster analysis (HCA) and principal component analysis (PCA) were performed using the SRplot web server (http://www.bioinformatics.com.cn/).

## 3 Results

### 3.1 Identification of Differentially Modified Peptides

Using a label-free quantitative proteome method, experimental data were obtained by LC-MS/MS analysis. A search based on open-pFind yielded detailed information on the mass spectra number of modified peptides in each sample, including the proteins they were located in and the types of modifications contained in the peptides. Modified peptides with reproducibility greater than 50% were screened separately in the elevated HbA1c group and the healthy control group, and the union of the two sets was used for subsequent analysis.

In the Group A comparison, a total of 6 833 modified peptides were identified. Based on the criteria of FC ≥ 1.5 or ≤ 0.67 and *p* < 0.05 in a two-tailed unpaired *t*-test analysis, 1 954 differentially modified peptides were identified in the mildly elevated HbA1c group compared with the healthy control group. Among these, 942 peptides exhibited a change from presence to absence, meaning that they were identified in more than half of the samples in the healthy control group but were not detected in any samples in the mildly elevated HbA1c group. In addition, 11 peptides exhibited a change from absence to presence, indicating that they were identified in more than half of the samples in the mildly elevated HbA1c group but were not detected in any samples in the healthy control group. Overall, 48.8% of the differentially modified peptides exhibited either a change from presence to absence or from absence to presence. Detailed information of all differentially modified peptides is listed in Table S1, including peptide sequences, modification types, and the proteins containing these peptides.

HCA and PCA were performed on total modified peptides, both of which distinguished the samples from the mildly elevated HbA1c group and the healthy control group (Figure 1).

**Figure 1.**
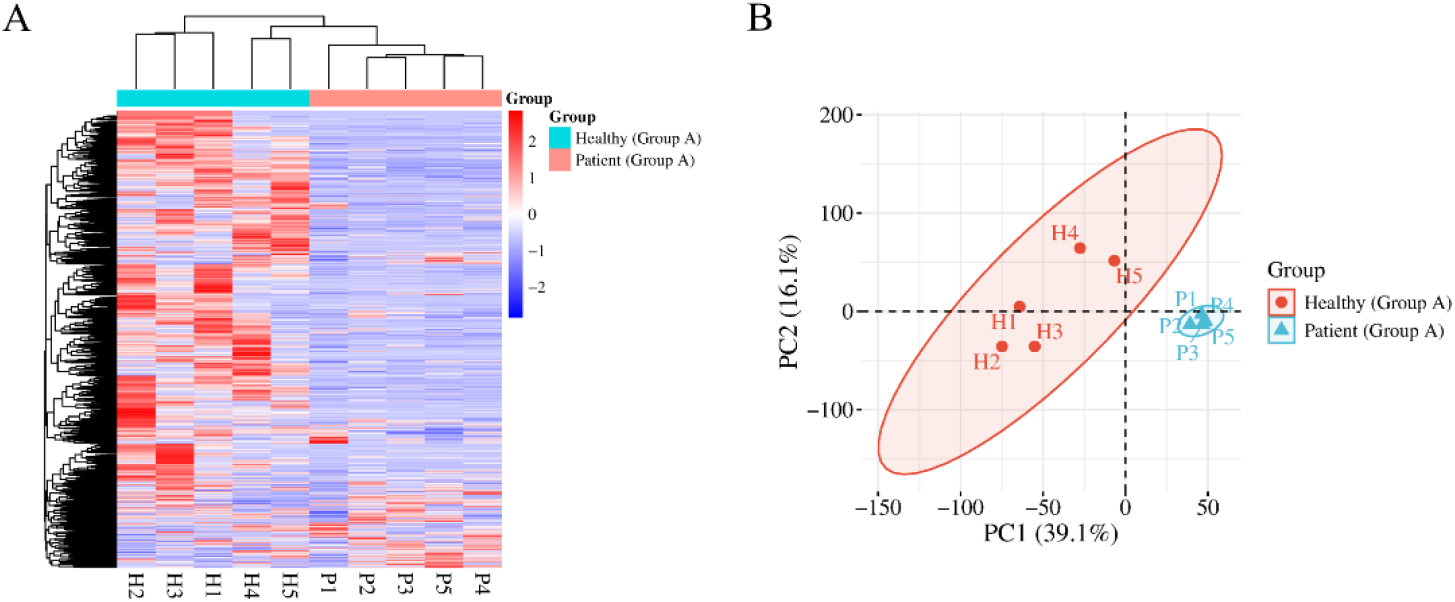
HCA and PCA of total modified peptides identified in the Group A comparison distinguishing samples from the mildly elevated HbA1c group and the healthy control group: (A) HCA; (B) PCA.

In the Group B comparison, a total of 8 162 modified peptides were identified. Using the same screening criteria (FC ≥ 1.5 or ≤ 0.67 and *p* < 0.05 in a two-tailed unpaired *t*-test analysis), 5 545 differentially modified peptides were identified in the elevated HbA1c group compared with the healthy control group. Among these, 778 peptides exhibited a change from presence to absence, whereas 4 017 peptides exhibited a change from absence to presence. Overall, 86.5% of the differentially modified peptides exhibited either a change from presence to absence or from absence to presence. Detailed information of all differentially modified peptides is listed in Table S2.

HCA and PCA were performed on total modified peptides, both of which distinguished the samples from the elevated HbA1c group and the healthy control group (Figure 2).

**Figure 2.**
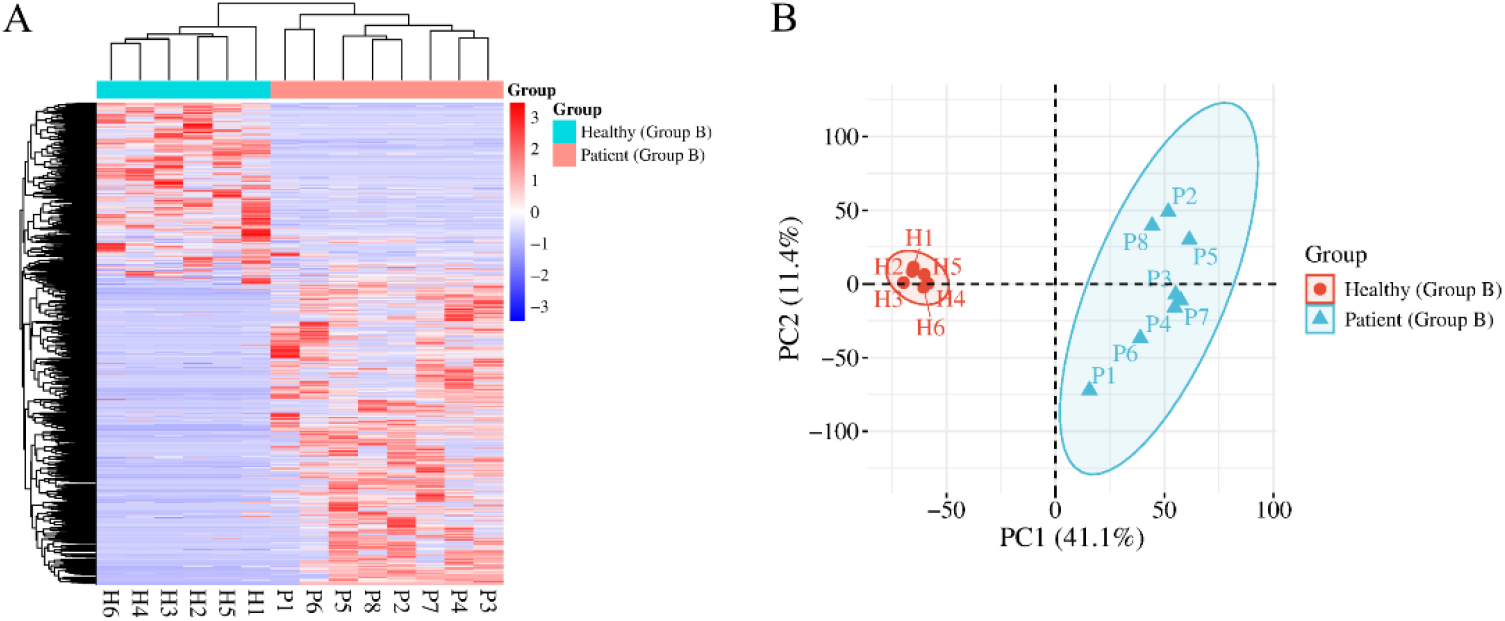
HCA and PCA of total modified peptides identified in the Group B comparison distinguishing the samples from the elevated HbA1c group and the healthy control group: (A) HCA; (B) PCA.

To assess the possibility of random generation of the identified differentially modified peptides, a randomized grouping test was performed on the total modified peptides in both comparisons. For Group A, 10 samples were randomly divided into two new groups, resulting in a total of 126 combinations. These combinations were then screened for differences based on the same criteria (FC ≥ 1.5 or ≤ 0.67, *p* < 0.05). The average number of differentially modified peptides yielded was 183.7, indicating that at least 90.6% of the differentially modified peptides identified in Group A were not randomly generated. For Group B, 14 samples were randomly divided into two new groups, resulting in a total of 3 003 combinations. Using the same screening criteria, the average number of differentially modified peptides yielded was 327.2, indicating that at least 94.1% of the differentially modified peptides identified in Group B were not randomly generated.

### 3.2 Analysis of Differentially Modified Peptides Commonly Identified by Both Groups

A total of 602 differentially modified peptides were commonly identified in both Group A and Group B, which may reflect the common changes associated with elevated HbA1c levels. Among these, 15 differentially modified peptides exhibited consistent and significant changes from presence to absence or vice versa in both groups (Table 2). Detailed information on all commonly identified differentially modified peptides is listed in Table S3, including peptide sequences, modification types, and the proteins containing these peptides.

**Table 2.**
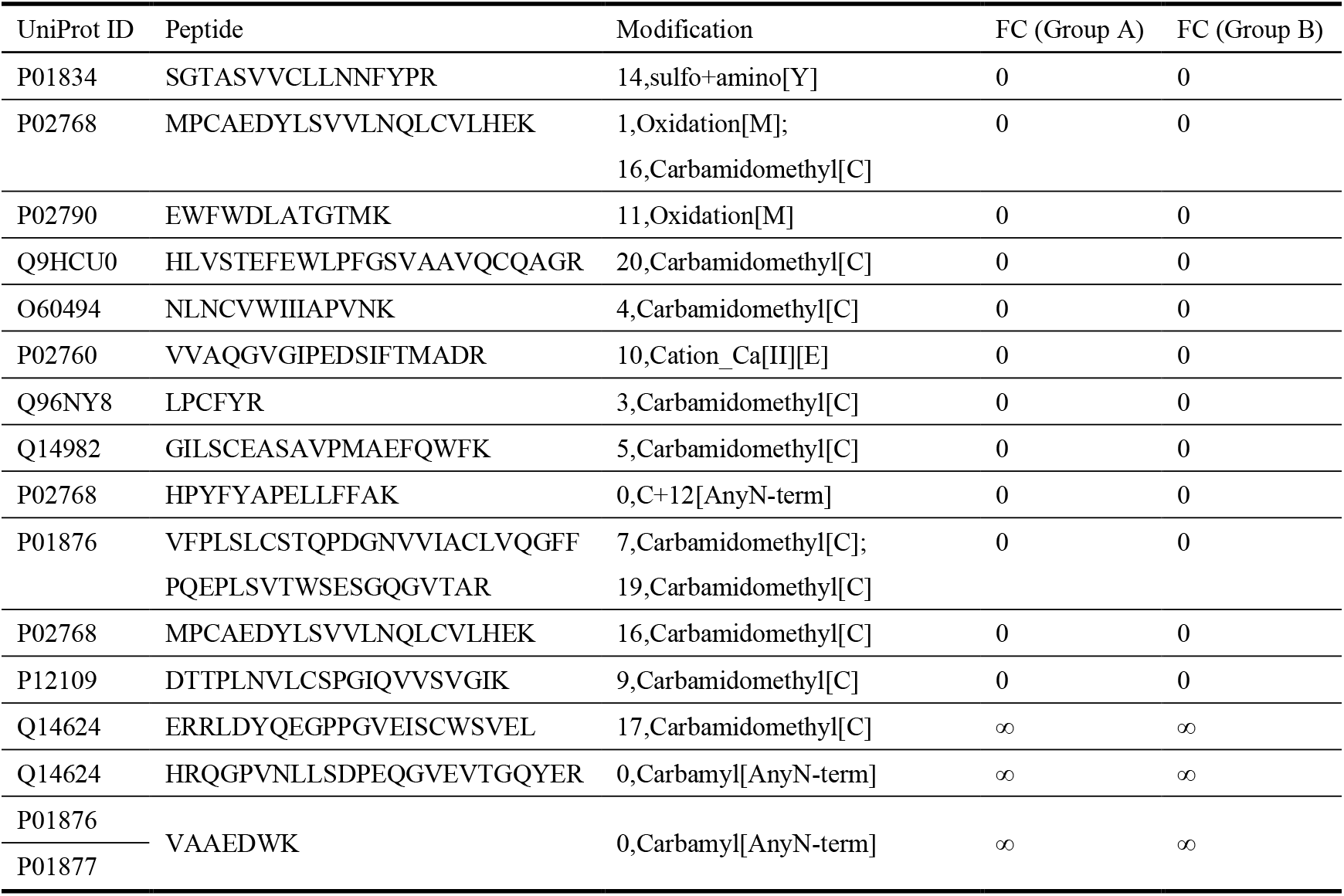
Differentially modified peptides exhibiting consistent changes from presence to absence or vice versa in both groups.

### 3.3 Analysis of Differential Modifications

The types of modifications and their identification frequencies in the differentially modified peptides from Group A and Group B were counted separately. A total of 160 modification types were identified in Group A, whereas 284 modification types were identified in Group B. The number of modification types increased with increasing HbA1c levels. Detailed information is listed in Table S4. The top 11 modification types ranked by identification frequency in both groups are shown in Table 3.

**Table 3.** Top 11 differential modification types by identification frequency.

| Group | Modification | Number | Group | Modification | Number |
| --- | --- | --- | --- | --- | --- |
| Group A | Carbamidomethyl[C] | 1206 | Group B | Carbamidomethyl[C] | 3084 |
|  | Oxidation[M] | 452 |  | Carbamyl[AnyN-term] | 1352 |
|  | Carbamyl[AnyN-term] | 241 |  | Deamidated[N] | 500 |
|  | Deamidated[N] | 154 |  | AEBS[Y] | 359 |
|  | GG[C](Dicarbamidomethyl[C]) | 137 |  | Oxidation[M] | 339 |
|  | Gln->pyro-Glu[AnyN-termQ] | 81 |  | GG[C](Dicarbamidomethyl[C]) | 239 |
|  | Pyro-carbamidomethyl[AnyN-termC] | 26 |  | Gln->pyro-Glu[AnyN-termQ] | 158 |
|  | Propionamide_2H(3)[C] | 23 |  | Oxidation[Y] | 157 |
|  | Dehydrated[D] | 21 |  | C+12[AnyN-term] | 65 |
|  | Cation_Ca[II][E] | 15 |  | AEBS[K] | 59 |
|  | Trp->Kynurenin[W] | 15 |  | Pyro-carbamidomethyl[AnyN-termC] | 58 |

In both groups, carbamidomethyl is the most frequent modification type, which is induced by the alkylation reagent iodoacetamide (IAA) acting on cysteine residues (Cys, C). This is an artificially introduced modification. It accounted for 43.7% of all identified differential modification types in Group A and 40% in Group B. In this study, samples from the mildly elevated HbA1c group and the healthy control group in Group A were processed using the same experimental procedures. No differences should have been observed if the state of the cysteine residues in both groups had been identical prior to treatment. However, the results showed that this modification was identified as significantly different between the two groups with an extremely high frequency. This suggests a pre-existing difference in the state of cysteine residues between the two groups before sample processing.

The sulfhydryl groups of cysteine residues form disulfide bonds through oxidation reactions, which are essential for maintaining protein stability [11]. Previous studies have shown that under diabetic conditions, protein disulfide isomerase (PDI) predominantly exists in a reduced form, rendering it incapable of forming disulfide bonds in newly synthesized proteins[12].

## 4 Discussion

HbA1c, as a biomarker of glycemic management [13], exhibits a continuous relationship with the risk of developing diabetes. In this study, mass spectrometry data from three published studies were integrated. Two patient groups with different HbA1c levels ([6.4±0.7]% and [8.6±1.6]%) were each compared with the healthy control group to systematically explore differences associated with different HbA1c levels from the perspective of urinary proteome modifications.

The results showed that urinary proteome modifications in both comparisons reflected changes associated with elevated HbA1c levels. A total of 1 954 differentially modified peptides were identified in the Group A comparison, and 5 545 were identified in the Group B comparison. Furthermore, the proportions of differentially modified peptides exhibiting changes from presence to absence or vice versa were 48.8% and 86.5%, respectively. Both the total number of differentially modified peptides and the proportion of differentially modified peptides showing significant presence-absence changes increased markedly with increasing HbA1c levels. Randomized grouping tests further validated the reliability of the experimental findings, indicating that at least 90.6% and 94.1% of the differentially modified peptides in each group were not randomly generated.

Taken together, the results from both comparisons indicate that different HbA1c levels are associated with distinct urinary proteome modification profiles, providing a basis for reflecting HbA1c levels through urinary proteome modifications. This study is a retrospective analysis with a relatively limited sample size, and age differences between the healthy control group and the elevated HbA1c group may have introduced confounding effects. Further validation can be conducted through large-scale clinical studies in the future.

## 5 Conclusion

In this study, we used urinary proteome modifications to investigate differences between two groups of patients with different HbA1c levels and healthy controls. Urinary proteome modifications comprehensively and systematically reflect changes associated with elevated HbA1c levels, with distinct modification profiles corresponding to different HbA1c levels. These findings suggest that urinary proteome modifications have the potential to reflect HbA1c levels and offer a new perspective for research on the early diagnosis of diabetes.

